# CRISPR Knockdown of *Kcnq3* Attenuates the M-current in NPY/AgRP Neurons

**DOI:** 10.1101/2020.05.28.122341

**Authors:** Todd L. Stincic, Martha A. Bosch, Avery C. Hunker, Barbara Juarez, Ashley M. Connors, Larry S. Zweifel, Oline K. Rønnekleiv, Martin J. Kelly

**Affiliations:** Department of Chemical Physiology and Biochemistry, Oregon Health and Science University, Portland, OR 97239; Division of Neuroscience, Oregon National Primate Research Center, Beaverton, OR 97006; Department of Psychiatry and Behavioral Sciences, University of Washington, 98195; Department of Pharmacology, University of Washington, Seattle, WA 98195

**Keywords:** Neuropeptide Y, fasting, 17β-estradiol, KCNQ channels, XE991

## Abstract

Arcuate nucleus Neuropeptide Y/Agouti-related peptide (NPY/AgRP) neurons drive ingestive behavior in response to the internal and external environment of an organism. NPY/AgRP neurons are adjacent to the median eminence, a circumventricular organ, and circulating metabolic factors and hormones communicate the energy state of the animal via these neurons by altering the excitability of NPY/AgRP neurons, which produces an appropriate change in behavior to maintain homeostasis. One example of this plasticity is seen in the M-current, a subthreshold, non-inactivating K^+^ current that acts to modulate excitability. Fasting decreases while estradiol increases the M-current through regulation of subunit mRNA expression of *Kcnq 2, 3, & 5*. KCNQ2/3 heteromers are thought to mediate the majority of the M-current. Here we used a recently developed single adeno-associated viral (AAV) vector containing a recombinase-dependent Staphylococcus aureus Cas9 (SaCas9) and a single guide RNA against *Kcnq3* to selectively delete *Kcnq3* in NPY/AgRP neurons to produce a loss of function in the M-current. We found that this virus was effective at knocking down *Kcnq3* but not *Kcnq2* expression. With the reduced KCNQ3 channel expression NPY/AgRP neurons were more depolarized, exhibited a higher input resistance, and the rheobase current needed to induce firing was significantly reduced, indicative of increased excitability. Although the resulting decrease in the M-current did not overtly alter ingestive behavior, it did significantly reduce the locomotor activity as measured in open field testing. Therefore, the SaCas9-sgKcnq3 is efficient to knock down *Kcnq3* expression thereby reducing the M-current and increasing the excitability of NPY/AgRP neurons.

## Introduction

In the hypothalamic arcuate nucleus (ARH), two distinct, neuronal cell types differentially modulate energy homeostasis – the proopiomelanocortin (POMC) and neuropeptide Y/agouti-related peptide (NPY/AgRP) neurons. NPY and AgRP are co-expressed in the same ARH neurons, and AgRP is the endogenous antagonist for the melanocortin (MC4) receptor, thus blocking the anorectic effects of the POMC peptide, α-melanocyte stimulating hormone [1, 2]. Ablation of NPY/AgRP neurons in adults causes rapid starvation [3, 4], while injection of NPY into the paraventricular nucleus stimulates food intake [5]. Optogenetic and chemo-genetic stimulation of NPY/AgRP neurons swiftly increases food consumption [6, 7], whereas stimulation of POMC neurons attenuates food intake [6, 8]. *In vivo* measurements of calcium activity have shown that NPY/AgRP neurons integrate anticipatory sensory cues to generate appropriate feeding behavior [9]. In fact, plasticity and responsiveness of ARH circuits are critical for the maintenance of energy balance.

In their orexigenic role, NPY neurons respond to many peripheral and central hormonal signals that control energy homeostasis [10–14]. NPY/AgRP and POMC neurons are also the major CNS targets of the anorectic hormone 17β-estradiol (E2) [15, 16]. E2 suppresses NPY expression in the ARH [17, 18]. Conversely, NPY/AgRP neurons are more excitable after fasting and leptin reduces their excitability [13]. Leptin and other peripheral signals *(e.g.* ghrelin and gastrin-releasing peptide) modulate NPY/AgRP excitability by targeting calcium and potassium (K^+^) channels [14, 19–21]. One such conductance, the M-current, is a sub-threshold, non-inactivating, voltage-dependent, outward K^+^ current. This K^+^ conductance modulates neuronal excitability, action potential kinetics and burst firing [22, 23]. The M-current is itself regulated by numerous neurotransmitters and neuropeptides that modulate neuronal excitability through G-protein coupled receptors including acetylcholine, serotonin, substance P and GnRH [22, 24].

The M-current amplitude is mediated by KCNQ channels [25, 26]. NPY/AgRP neurons highly express the transcripts for the subunits KCNQ 2, 3, and 5, but not 4, with most individual cells expressing detectable levels of two or more of these subunits [27] to form hetero-multimeric [28], and possibly homo-multimeric complexes [29]. Global knockout of *Kcnq2* is perinatally lethal [30] unlike *Kcnq3* [29] or *Kcnq5* [31]. When *Kcnq2 or Kcnq3* are individually expressed in *Xenopus* oocytes the M-current is negligible, but equimolar injections of the transcripts produce a significantly greater current [25], suggesting that these heteromers contribute the majority of the native conductance or that KCNQ3 coexpression aids in trafficking to the membrane [32–34]. However, these assumptions have been called into question using conditional knockout mice. When KCNQ subunits are individually deleted from cortical pyramidal cells, only *Kcnq2* knockout produces a robust increase in neuronal excitability, suggesting that KCNQ2 is obligatory [35]. Currently, it is unknown whether this represents a general rule for KCNQ channels or is cell-type specific. The role and importance of KCNQ5 in the M-current is not as well characterized. KCNQ5 subunits are thought to exist in homomeric or KCNQ3/5 heteromeric complexes [36, 37]. In the hippocampus, expression of *Kcnq5* has been suggested to decrease the M-current due to the competition for KCNQ3 subunits to the detriment of KCNQ2/3 heteromer formation [31]. In the ARH, however, elevated *Kcnq5* expression appears to contribute to the E2-potentiation of the M-current [27]. Surprisingly, *Kcnq2* KO mice also have a 40% loss of KCNQ5 protein, though the underlying cause remains unknown [35]. Previous KCNQ3 knockouts have been global [29–31] or present during development [35, 38] and, therefore, the influence of compensatory mechanisms cannot be dismissed. The newly developed Cre-dependent SaCas9 vectors enable region and cell-type specificity in an adult animal. Using this novel approach we tested the hypothesis that knockout (KO)/knockdown (KD) of *Kcnq3* in NPY/AgRP neurons in adult animals would increase their excitability, perhaps leading to disruptions in the regulation of energy balance.

## Materials & Methods

### Animals and Treatments

All animal procedures described in this study were performed in accordance with institutional guidelines based on National Institutes of Health standards and approved by the Institutional Animal Care and Use Committee at Oregon Health and Science University.

### Mice

Npy^hrGFP^ (JAX #006417) [14] or AgRP^Cre^ (JAX #012899) [39] transgenic mice (breeders provided by Dr. Brad Lowell, Harvard University, Cambridge, MA, USA) were selectively bred in-house and maintained under controlled temperature (25°C) and photoperiod conditions (lights on at 6 a.m. and off at 6 p.m.) with food (5L0D; Laboratory Diets, St. Louis, MO, USA) and water ad libitum unless otherwise stated. For single-cell reverse transcription polymerase chain reaction (scRT-PCR) and electrophysiology only female mice were only used, whereas behavior was measured solely in males.

### Visualized Whole-Cell Patch Recordings

Coronal brain slices (240 μm) containing the ARH from gonadectomized female or intact male mice were made in an ice-cold sucrose cutting solution (see recipe below) and stored in a bubbled (95% 0_2_ & 5% CO_2_) chamber containing artificial cerebrospinal fluid (aCSF; see recipe below). Whole-cell patch recordings were made from eGFP/YFP/mCh fluorescent neurons using an Olympus BX51W1 upright microscope with an Exfo X-Cite 120 Series fluorescence light source, epifluorescence (FITC and mCherry filters) and video-enhanced, infrared-differential interference contrast. Patch pipettes were filled with K^+^-gluconate internal solution (in mM): 128 potassium gluconate, 10 NaCl, 1 MgCl_2_, 11 EGTA, 10 HEPES, 3 ATP, and 0.25 GTP (pH was adjusted to 7.3–7.4 with 1N KOH, 290–300 mOsm). Pipette resistances ranged from 3-5 MΩ. in whole cell configuration. Cells were excluded from analysis if their input resistance was below 800 MΩ and if access resistance remained less than 20 MΩ throughout the experiment. I-V relationships of currents were examined with voltage steps.

### Electrophysiology data analysis

Electrophysiological signals were digitized with Digidata 1440A (Molecular Devices), amplified with an Axopatch 200B (Molecular Devices), and analyzed using ClampFit version 10 software (RRID:SCR_011323, v10.3, Molecular Devices). Subsequently, all electrophysiology data was transferred to Sigma Plot 13 (Jandel Scientific, San Rafael, CA) or Graph Pad Prism 6/7 (La Jolla, CA) for statistical analysis. The liquid junction potential was corrected for all data analysis. All values are expressed as Mean ± SEM. Comparisons between two groups were made using un-paired Student’s t-test or between multiple groups using an ANOVA (with post hoc comparisons) with p-values < 0.05 considered significant. When variances differed significantly, Mann-Whitney was used instead.

### Electrophysiological Solutions and Drugs

A sucrose solution was used during Vibratome slicing (in mM): 2 KCl, 1 MgCl_2_-6H_2_O, NaH_2_PO_4_, 10 HEPES, 10 glucose, 208 sucrose, 26 NaHCO_3_, 2 MgSO_4_-7H_2_O, and 1 CaCl_2_. For harvesting and electrophysiological recordings, a standard artificial cerebrospinal fluid was used (in mM): 124 NaCl, 5 KCl, 1.44 NaH_2_PO_4_, 5 HEPES, 10 glucose, 26 NaHCO_3_, 2 MgSO_4_-7H_2_O, and 2 CaCl_2_. Tetrodotoxin (1 μM, Alomone Labs, Jerusalem, Israel) was added to the bath for all recordings except when measuring the rheobase.

### AAV Delivery

Viruses were prepared at the University of Washington according to published methods [40, 41]. Four to eight weeks prior to each experiment, AgRP^Cre^ mice (>60 days old) received bilateral ARH injections of Cre-dependent adeno-associated (AAV; serotype 1) viral vectors encoding yellow fluorescent protein, YFP (AAV-EF1α-YFP), or mCherry, mCh (AAV-EF1α-mCh), either alone or co-injected with an AAV1 designed to encode SaCas9 and a single guide RNA (sgRNA). See SaCas9 section for specifics on sgRNA design. Using aseptic technique, anesthetized (1-1.5% isoflurane/O_2_) mice received a medial skin incision to expose the surface of the skull. The glass pipette (#3-000-203-G/X; Drummond Scientific, Broomall, PA) with a beveled tip (diameter = 45 μm) was filled with mineral oil, loaded with an aliquot of AAV using a Nanoject II (Drummond Scientific). ARH injection coordinates were anteroposterior (AP): −1.15 mm, mediolateral (ML): ± 0.33 mm, dorsoventral (DV): −5.80 and −5.70 (surface of the brain z = 0.0 mm); 250 nl of the AAV (2 × 10^12^ particles/ml) were injected (100 nl/min) at each position, the pipette was left in place for 10 min post-injection and then slowly retracted from the brain. The skin incision was closed using Vetbond (3M) and each mouse received analgesia (Rimadyl; 4-5 mg/kg, sc).

### Gonadectomy

Male gonads were left intact, and all females were subjected to ovariectomy (OVX) at least 7 days prior to each experiment. Rimadyl (4-5 mg/kg, *sc*) was given immediately after surgery for relief of postoperative pain. For the quantitative measurement of CRISPR gene editing, we used fed, OVX female mice that were treated with 17β-estradiol benzoate (EB). We selected this model in females to enhance expression of *Kcnq 2, 3 and 5 [27]* with the goal of maximizing the sensitivity to detect reductions in transcripts and currents. Females received an injection of a low (0.25 μg) and then a high (1.5 μg) dose of E2 benzoate administered in the morning of the two days preceding experiments. We have documented that this regimen induces a pre-ovulatory surge of luteinizing hormone in females, replicating proestrus [42, 43]. Circulating levels of E2 were verified by the uterine weights (>95 mg) at the time of hypothalamic slice preparation (between 8:30 & 10:30 am).

### Generation and validation of AAV1-FLEX-SaCas9-U6-sgKcnq3

The sgRNA for targeting the *Kcnq3* locus was designed as previously described [41]. The following oligos (Sigma) were used to clone into pAAV-FLEX-SaCas9-U6-sgRNA (Addgene 124844). *Kcnq3* forward: CACCGCCACCGCCGAGTCCCCTCCAG; *Kcnq3* reverse: AAACCTGGAGGGGACTCGGCGGTGGC.

Targeted deep sequencing of *Kcnq3* locus: AAV1-FLEX-SaCas9-U6-sg*Kcnq3* and AAV1-FLEX-EGFP-KASH were co-injected into the VTA of DAT-Cre mice. Nuclei were isolated and FACS for EGFP-KASH-positive and -negative nuclei was performed, followed by targeted deep sequencing of whole-genome amplified (WGA) DNA as described [41]. Four weeks after surgery, tissue punches of the ventral midbrain were pooled from 3 mice into a single group and homogenized in 2mL of homogenization buffer containing (in mM): 320 Sucrose (sterile filtered), 5 CaCl_2_ (sterile filtered), 3 Mg(Ac)_2_ (sterile filtered), 10 Tris pH 7.8 (sterile filtered), 0.1 EDTA pH 8 (sterile filtered), 0.1% NP40, 0.1 Protease Inhibitor Cocktail (PIC, Sigma), 1 β-mercaptoethanol. Homogenization was performed using 2mL glass dounces (Sigma Cat#: D8938-1SET); 25 times with pestle A, then 25 times with pestle B. The volume of the homogenate was transferred to a 15mL conical tube and brought up to 5mL using homogenization buffer, mixed, and incubated on ice for 5 minutes. 5mL of 50% Optiprep density gradient medium (Sigma Cat#: D1556-250ML) containing (in mM): 5 CaCl_2_ (sterile filtered), 3 Mg(Ac)_2_ (sterile filtered), 10 Tris pH 7.8 (sterile filtered), 0.1 PIC, 1 β-mercaptoethanol was added to the homogenate and mixed by inversion. The suspension was layered onto 10mL of 29% iso-osmolar Optiprep solution in a 1×3 ½ in a Beckman centrifuge tube (SW32 Ti rotor) and centrifuged at 7500 RPM for 30min at 4°C. Floating cell debris was removed and the supernatant gently poured out. The pellet was resuspended in sterile 1xPBS. 500 GFP-positive and 500 GFP-negative nuclei were sorted directly into 3uL of REPLI-g Advanced Storage buffer (Qiagen Cat#: 150365) in a PCR tube strip (Genessee Cat#: 24-706) using a BD AriaFACS III. Whole genome amplification (WGA) was performed directly following FACS using the REPLI-g Advanced DNA Single Cell kit (Qiagen Cat#: 150365) according to manufacturer’s instructions. For generation of amplicons, 1ul of WGA DNA was diluted 1:50 and amplified (PCR 1) with Phusion High Fidelity Polymerase (Thermo Fisher Cat#: F530L) using the following thermocycler protocol: initial denaturation (30sec, 95°C); denaturation (10sec, 95°C); annealing (20sec, 69°C); extension (10sec, 72°C); cycle repeated ×34; final extension (5min, 72°C). For PCR 2, 1uL of PCR 1 was amplified with a second set of primers using the same thermocycler protocol. The 205bp amplicon from PCR 2 was gel extracted using the MinElute gel extraction kit (Qiagen Cat#: 28606), and sent to Genewiz for Amplicon-EZ targeted deep sequencing and Sanger sequencing. The primers used for the generation of the amplicons were as follows: PCR 1 forward: CAAGTGCTCCTACTTCCC reverse: GTCTTTGCCAGGAGCCCGAT, PCR 2 forward: AGATGGGTCTCAAGGCTC, reverse: AGGGTCCCGTCTTTGTCG.

### Single Cell Harvesting

Ovariectomized (OVX), EB-treated female mice were rapidly decapitated and coronal brain sections (240 μm) were cut using a Vibratome (VT-1000S; Leica, Wetzlar, Germany). The ARH was microdissected from basal hypothalamic slices (3-4 slices per mouse). Gentle trituration following incubation with papain (Sigma-Aldrich) was used to dissociate the ARH neurons before dispersion onto a glass bottom dish where the healthy cells settled and adhered to the glass bottom. After 15 minutes, the aCSF was removed and fresh artificial cerebrospinal fluid (aCSF) was added to the plate. This washing procedure was repeated two times. Throughout the dispersion and harvesting procedure, a constant flow (2 ml/min) of oxygenated aCSF circulated into the plate while the effluent circulated out using a peristaltic pump. The aCSF flow helped ensure fresh, oxygenated media was reaching the cells and assisted in clearing out unhealthy cells and debris from the trituration. The cells harvested were those observed to be fully intact, with one to three processes and a smooth cell membrane (**Fig 2A**) as visualized using an inverted microscope (DMIL; Leica) equipped with a fluorescent LED light source (X-Cite 110LED; Excelitas Technologies Corp., Waltham, MA). Individual neurons were patched, and then harvested with gentle suction into the pipette using a XenoWorks Micromanipulator/Microinjector system (Sutter Instrument Company, Novato, CA) and expelled into a siliconized 0.65 ml microcentrifuge tube containing 5X Superscript III buffer (Invitrogen, Carlsbad, CA), 15 U RNasin (Promega, Madison, WI), 10 mM dithiothreitol (DTT), and diethylpyrocarbonate (DEPC)-treated water in a total of 8 μl for quantitative real-time PCR (qPCR) (5 or 10-cell pool/tube). After electrophysiological experiments, the cytosol of recorded cells was harvested with gentle suction into the recording pipette for *post hoc* identification with scRT-PCR. Each single cell was expelled in a siliconized 0.65 ml microcentrifuge tube containing the solution described above and both cell pools and single cells were stored at −80 C prior to reverse transcription.

### Quantitative RT-PCR and single Cell RT-PCR and Primer design

cDNA synthesis was performed in a reaction volume of 25 μl (10 cell pools) or 20 μl (single cells) containing dNTPs (0.5 mM, Promega), random primers (100 ng/per tube, Promega), anchored oligo(dT)20 primers (400 ng/tube, Invitrogen), Superscript III reverse-transcriptase (100 U/per tube, Invitrogen), RNAsin (15 U), DTT (6 mM) and DEPC-treated water according to established protocols [42] and stored at −20 °C. Primers for the genes that encode for *Kcnq2*, *Kcnq3* and β-actin were designed to cross at least one intron-exon boundary and are listed as follows: *Kcnq2* (accession number NM_010611) 92 bp PCR product, forward primer 1079-1098 nt reverse primer 1151-1170 nt; *Kcnq3* (accession number NM_152923) 94 bp PCR product, forward primer 879-896 nt, reverse primer 955-972 nt; *Actb* (β-actin) (accession number NM_007393) 110 bp PCR product, forward primer 446-465 nt, reverse primer 535-555 nt. Of note, our first choice for primer location for Kcnq3 was to span the PAM and sgRNA sites. However, the PAM and sgRNA sites are found at nt 488-514 (exon1) of the 5’ end of the Kcnq3 mRNA sequence (NM_152923.3), an area rich in GC content (76%) making it essentially impossible to design adequate primers for scRT-PCR or qPCR analysis. Therefore, we designed a highly efficient set of primers noted above (Forward nt 879-896, reverse nt 885-886) producing a 94 bp product spanning the intron-exon boundary between exons 2 and 3.

qPCR was performed on a QuantStudio 7 Flex Real-Time PCR System (ThermoFisher Scientific) using the Power Sybr Green (ThermoFisher Scientific) mastermix method. The efficiencies of the target and reference gene amplifications was determined by serially diluting cDNAs (1:50 to 1:12,800) from mouse basal hypothalamus to construct standard curves. The efficiency was calculated per the following formula: E=10^(−1/m)^-1, m=slope [44, 45]. The comparative ΔΔC_⊤_ method was used to determine values from duplicate samples of 4 μl for the target genes (*Kcnq2*, *Kcnq3*) and 2 μl for the reference gene (*Actb*). The relative linear quantity was determined using the 2^−ΔΔCT^ equation [42]. In order to determine relative expression levels of mRNA in neurons obtained from control and KCNQ3 knockdown mice, the mean ΔCT for the target gene from the control samples was used as the calibrator, and the data expressed as an n-fold change in gene expression normalized to the reference gene, *Actb,* and relative to the calibrator. A two-tailed, unpaired t-test was used for analysis and data is expressed as mean ± SEM, * p<0.05, ** p<0.01, and *** p<0.001.

### Open field

Animals were tested during their relative light phase between ZT 6-8. At least an hour before testing to allow mice to acclimate in a dimly lit room adjacent to the testing apparatus in the presence of white noise. The open field test was conducted in 40×40 cm opaque white boxes cleaned with acetic acid at the beginning and end of testing day and with 70% ethanol between subjects, allowing time for any odor to dissipate. The room light was adjusted to 200 lux with white noise also present. Mice were tracked using Ethovision XT 10 (Noldus, Leesburg, VA) for 10 min. The distance (cm) traveled and average velocity (cm/s) was calculated in addition to percent time spent and frequency of crossing into the center, the latter two considered measures of anxiety.

### Lickometer

Med Associates operant chambers (ENV-307W) equipped with dual lickometers contained in sound attenuating cubicles (ENV-022V) were used to measure vanilla Ensure (Abbott Nutrition) consumption. Mice were given access to two sipper tubes, either for Ensure (36g in 100 ml tap water) or tap water. The house light was on throughout the testing period of sixty minutes. The day prior to start of training a small dish containing Ensure was placed in mouse home cages to prevent neophobia. Chambers were controlled and data was collected with MedPC 5 software (Med Associates) and exported to Excel before analysis in Graphpad Prism 6/7.

### Food intake measurements

Mice were singly housed with crinkle paper enrichment and ramekin bowls containing standard food pellets with ad libitum access to water. After two days of habituation, all food from the hopper was removed and 12-14 g of chow was placed in the ramekin. In the morning on the following two days the remaining pellets and crumbs were weighed, and the difference marked as baseline food consumed. The mice were then moved to clean cages (to prevent coprophagy) with water, but no food for an overnight fast. After 16 hours food was restored and food intake measured for an additional day (refeed) [46].

## Results

### CRISPR Kcnq3 significantly decreases Kcnq3 expression

The CRISPR/Cas9 system is recognized as an efficient means of generating insertion/deletion (indel) mutations to cause a loss of function with specific genes [47–49]. However, due to the large size of *Staphylococccus pyogenes*, there is insufficient capacity in the expression cassette of adeno-associated viruses to also include the single guide RNA (sgRNA) necessary for gene targeting. This limitation requires use of a separate viral vector [50, 51] or incorporation of a Cre-dependent SpCas9 transgenic line [52]. We recently developed a single viral vector for conditional expression of the smaller *Staphylococcus aureus* (SaCas9) and sgRNA that yields high-efficiency mutagenesis in specific cell types [41]. To inactivate *Kcnq3*, we generated a guide targeting exon 1 which is conserved across all splice variants. Due to the small size of the ARH and the relatively small number of AgRP neurons, we confirmed the efficacy of SaCas9 mutagenesis of *Kcnq3* in midbrain dopamine neurons, as described (Hunker et al., 2020). Briefly, DAT-Cre mice were co-injected with AAV1-FLEX-SaCas9-U6-sg*Kcnq3* and AAV1-FLEX-EGFP-KASH and tissue was harvested four weeks following surgery. EGFP-positive and negative nuclei were isolated via FACS and whole genome amplification (WGA) was performed followed by targeted deep sequencing of a PCR amplicon containing the targeted region of *Kcnq3* and the CRISPOR-predicted off-target *Grm8* (Figure 1A-C). SaCas9 generated numerous insertions and deletions (indels) centered at 3 base pairs upstream of the protospacer adjacent motif (PAM) (Figure 1A, D-E). A small number of base changes (2%) and deletions (1%) were observed in GFP-negative nuclei and similarly small number of base changes were observed in *Grm8*, likely reflecting the fidelity of sequencing (Figure 1B-C, E).

**Figure 1:**
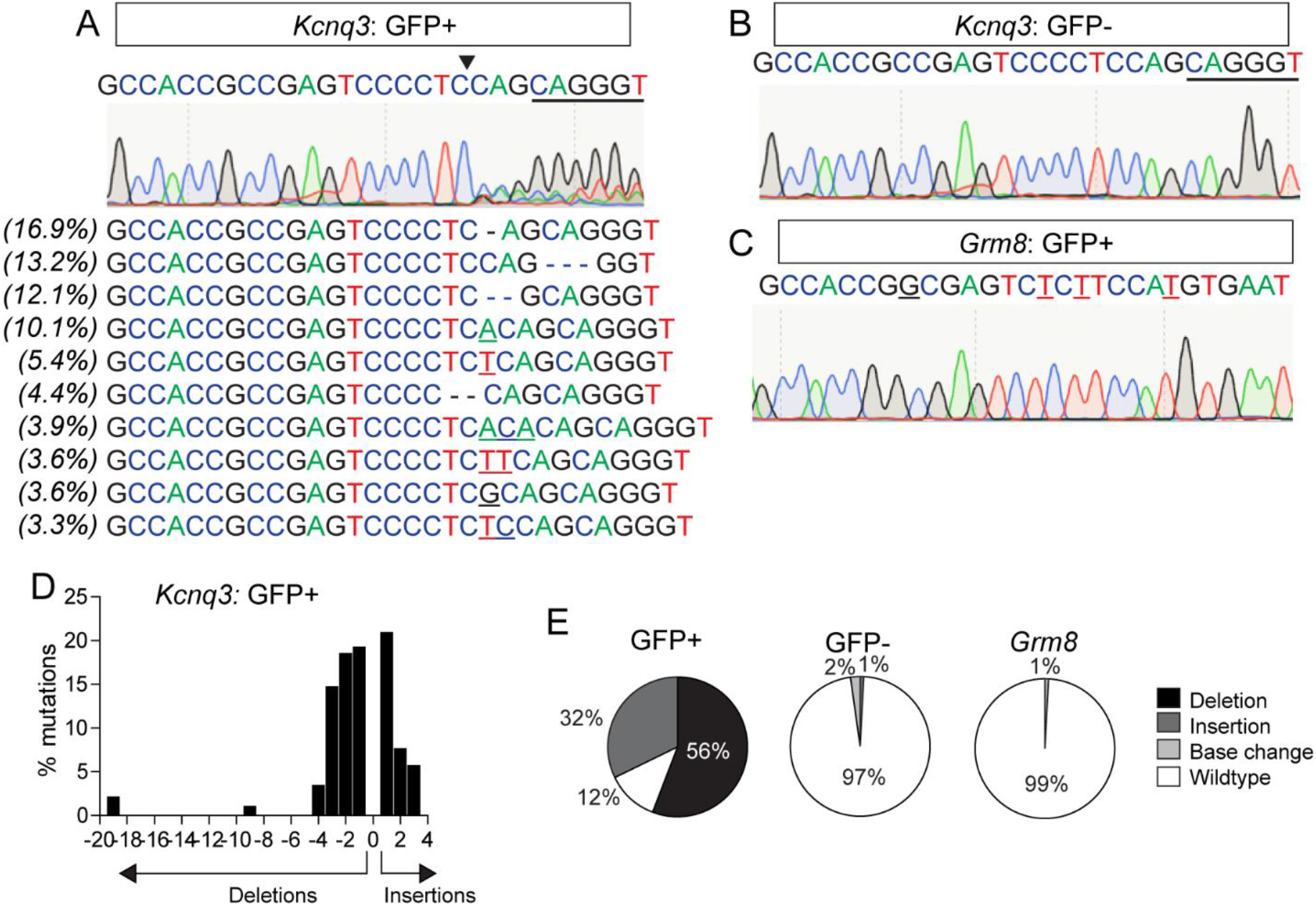
Sequencing of *Kcnq3* in sgKcnq3-targeted mice. A) Sequencing of GFP+ nuclei. Top: sgKcnq3 sequence with PAM underlined and SaCas9 cut site indicated by black arrow. Middle: Sanger sequencing results displaying multiple peaks beginning at the cut site. Bottom: Top ten mutations at cut site with the percent of total reads they occur on the left. Base changes: bolded. Insertions: underlined. Deletions: marked with a “−”. B) Sequencing of GFP-nuclei displaying no evidence of mutations at SaCas9 cut site. C) Sequencing of sgKcnq3 predicted off-target in exon of *Grm8* displaying no evidence of mutations at SaCas9 cut site. D) Frequency distribution of insertions and deletions in *Kcnq3* from GFP+ nuclei. E) Percent of wildtype, deletions, insertions, and base changes as percent of total reads for (left) *Kcnq3* in GFP+ and GFP-nuclei (right) the exon off-target *Grm8*.

**Figure 2:**
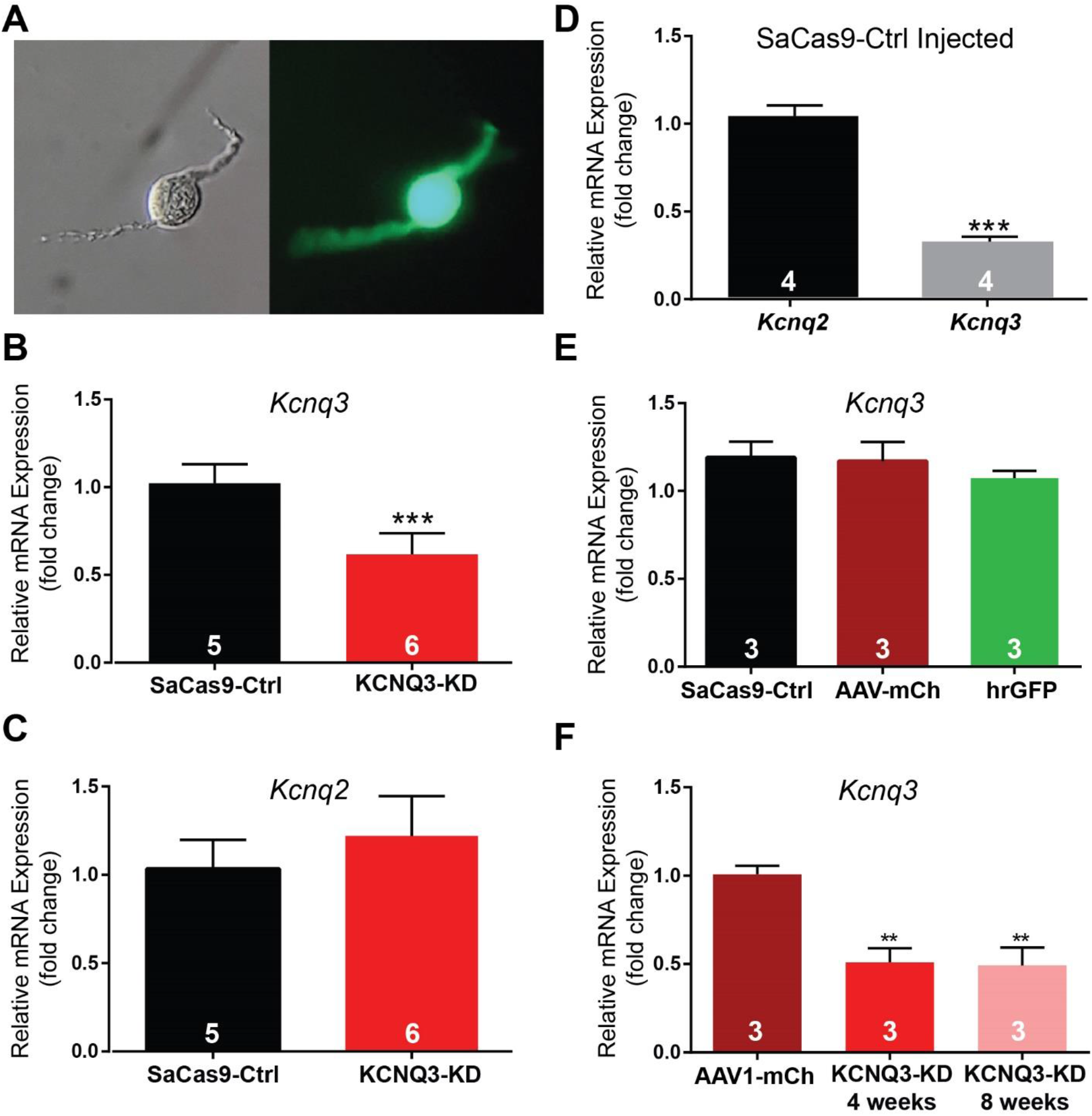
Quantitative RT-PCR on NPY/AgRP neurons. A) After digestion and dispersal, individual fluorescent cells were collected. Brightfield image of a dissociated cell with intact processes (left). Fluorescence was used to confirm cell type before harvesting (right). B) Relative expression of *Kcnq3* mRNA (10-cell pools, 4/animal), normalized to the control average (1.02±0.05, n=5) was significantly reduced in knockdown animals (0.62±0.05, n=6, p<0.001) using an unpaired *t*-test, *t*_(9)_ = 5.811. C) The relative expression of *Kcnq2* (10-cell pools, 4/animal), another KCNQ channel subunit, was unaffected by the control (1.03±0.07, n=5) or the targeted SaCas9 virus (1.22±0.23, n=6; unpaired t-test,*t*_(9)_ = 1.535 p>0.05). D) The *Kcnq2* transcript (1.03±0.06, n=4) is significantly more abundant than *Kcnq3* (0.32±0.03, n=4; unpaired *t*-test, *t*_(6)_ = 10.52). Only data from control animals is shown (10-cell pools, 4/animal). E) The relative expression of *Kcnq3* mRNA (5-cell pools/ 4/animal) was no different between several control groups which included AgRP^Cre^ animals injected with the SaCas9-Control (0.97±0.05) and AAV-DIO-mCh, AAV-DIO-mCh alone (0.91±0.05), or uninjected NPY^hrGFP^ mice (0.97±0.07) using a One-way ANOVA (F_(2,6)_ = 0.195, p>0.05). F) Knockdown of *Kcnq3* (5-cell pools, 4/animal, n=3/group) is stable between 4-weeks (0.49±0.1, unpaired *t*-test, *t*_(4)_ = 5.370) and 8-weeks (0.51±0.14, unpaired *t*-test, *t_(4_)* = 4.626) post-injection compared to 4-week fluorophore controls (1.01±0.05). ** p<0.01 and*** p<0.001. Error bars indicate SEM.

Based on these promising results of *Kcnq3* editing by the SaCas9 vector, a cohort of females were given bilateral stereotaxic injections in the ARH of either AAV1-FLEX-SaCas9-sg*Kcnq3* or KCNQ3 knockdown (KCNQ3-KD) virus or a control virus containing the *Kcnq3* guide with three base pairs in the seed region mutated (SaCas9-control) as described [41]. An additional Cre-dependent virus of the same serotype (AAV1) that drove expression of a fluorophore (YFP or mCh) was co-administered in order to visualize injection quality and facilitate harvesting of cells (**Fig 2A**). After three weeks, mice underwent OVX and were subsequently given EB-treatment on the two days prior to sacrifice. Brain slices were prepared, and cells harvested as previously described [53] and analyzed with qPCR. We found that the KCNQ3-KD group displayed a reduction in relative expression of *Kcnq3* in NPY/AgRP neurons compared to the control group (**Fig 2B**), but *Kcnq2* mRNA was not changed in NPY/AgRP neurons in response to *Kcnq3* KD (**Fig 2C**). Of note, the relative expression of *Kcnq2 mRNA* was much more abundant than *Kcnq3 mRNA* (**Fig. 2D**). Hence, the qPCR data corroborated the Kcnq3 sequencing data in the sgKcnq3-targeted mice (**Fig. 1A**) that we have selectively reduced *Kcnq3* gene expression in NPY/AgRP neurons.

Next, we tested whether the procedure itself affected *Kcnq3* mRNA expression compared to the control SaCas9 virus. The NPY^hrGFP^ and AgRP^Cre^ lines have both been bred to congenicity with C57BL/6J, and in the ARH, NPY and AgRP neurons are functionally identical. Therefore, we expected their relative expression of *Kcnq3* to be similar. However, the Cre-expression, stereotactic surgery, or viral infection could have affected cell health and transcription. Therefore, AgRP^Cre^ female mice were given ARH injections of AAV1-EF1α-DIO-mCh and after three weeks, mice underwent OVX and were subsequently given EB-treatment on the two days prior to sacrifice. After 4 weeks post-injections animals were sacrificed, and tissue processed for qPCR as described above. Age-matched NPY^hrGFP^ female mice were also selected and processed. As expected, there was no significant difference between any of the groups (**Fig 2E**). Together these data indicate that the observed reduction in *Kcnq3* mRNA expression is solely due to specific CRISPR gene editing by the vector with the correctly targeted sgRNA.

Finally, we wanted to determine the extent and stability of the knockdown. This is an important detail to develop a time frame in which to conduct behavioral and electrophysiological experiments. Therefore, six female AgRP^Cre^ mice were once again prepared as described above. At 4-5 weeks, three of the mice were euthanized and fluorescent ARH cells harvested. The remaining three mice were processed 8-10 weeks after injection. *Kcnq3* relative expression was reduced at both the 4-week and 8-week marks compared to mice injected with only the fluorophore virus and there was no difference between the two time points (**Fig 2F**). This data would indicate that the full effect of the single vector CRISPR gene editing is achieved by four weeks and remains stable, at least within this window of time, and allows ample time to conduct experiments testing the effects of KCNQ3 knockdown.

### Selective deletion of Kcnq3 reduces the M-current in NPY neurons

The M-current is a sub-threshold K^+^ current that influences the resting membrane potential to regulate the firing of a neuron. Fasting and loss of E2 reduce the amplitude of the M-current in NPY/AgRP neurons, increasing their excitability [27] and presumably underlying, in part, the hyperphagia associated with those conditions. Therefore, we initially measured the effects of CRISPR-mediated deletion of *Kcnq3* in NPY/AgRP neurons from fed, EB-treated OVX females. Most patch clamp recordings were taken from slices adjacent to those used for qPCR experiments, enabling gene transcription and electrophysiology data to be collected from the same animal. For NPY/AgRP cells while there was a significantly higher input resistance in KD cells (Ctrl 1.2±0.1 GΩ, n=31 & KCNQ3-KD 1.6±.0.1 GΩ, n=54; Unpaired *t*-test, *t_(50)_* = 3.358; p<0.01), the membrane capacitance (Ctrl 13.9±0.6 pF & KCNQ3-KD 14.7±0.5 pF; Unpaired *t*-test, *t*(_67)_ = 0.9767, p>0.05) and resting membrane potential (Ctrl: −50.8±1.7 mV, n=31 & KCNQ3-KD: −47.4±1.6 mV, n=54; Unpaired *t*-test, *t*_(83)_ = 1.349, p>0.05) were no different.

The M-current contributes to whole-cell K^+^ currents and can be elucidated in voltage clamp through established protocols [24, 54]. The deactivation or relaxation was measured as the difference between the steady state (>400 ms) and instantaneous (~5 ms after voltage step) current (**Fig 3A**). The membrane potential was stepped from −60 (V_hold_) to −20 mV to open the maximal number of KCNQ channels before stepping down to more hyperpolarized voltages (−25 to −75 mV). The current relaxation represents the deactivation of the KCNQ channels with greater “sags” indicating that a larger M-current is present [27]. Current responses to this protocol were measured in the presence of tetrodotoxin (1 μM, TTX) to prevent sodium driven action potentials (**Fig 3 B & C**). In the KCNQ3 KD group the IV plot shifted rightwards, and the M-current was significantly smaller at more depolarized voltages (**Fig 3D**, −45 to −25 mV). As this is a subthreshold current, the amplitude of the M-current should be greater in this range of membrane potentials [23], which would further support the selective nature of the CRISPR gene editing. XE 991 (40 μm) is a selective and potent blocker of KCNQ channels that quickly inhibits the M-current [27] (**Fig 3E**). XE displayed equivalent efficacy blocking the M-current in control and KCNQ3-KD animals (**Fig 3F**), which is also seen when recording from hippocampal neurons in KCNQ3 KO mice [29].

**Figure 3.**
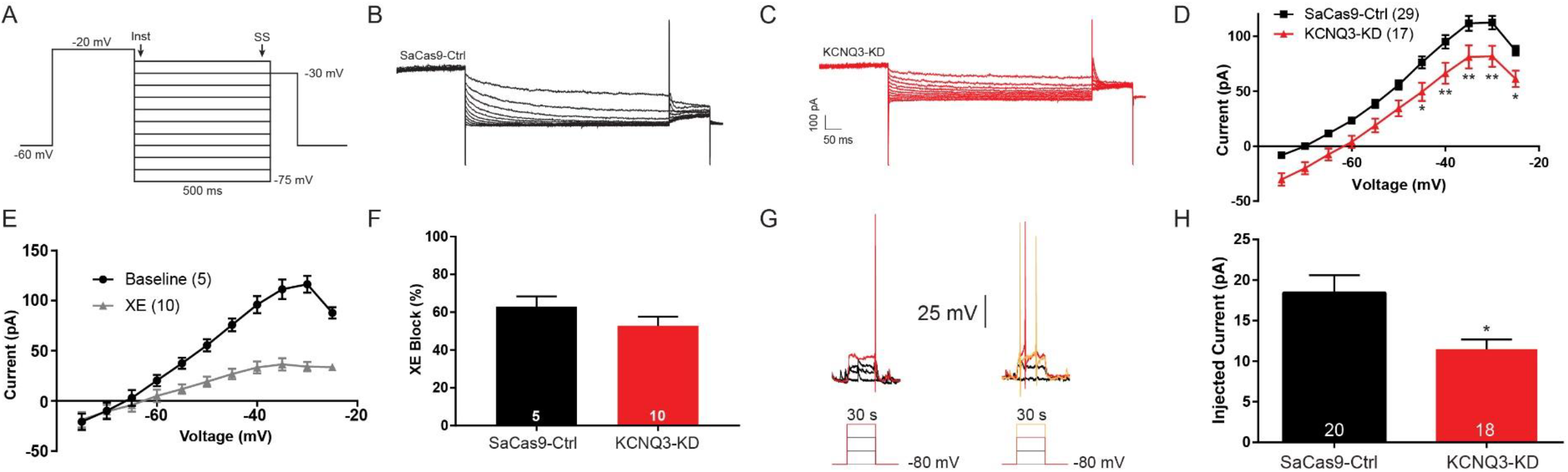
CRISPR gene editing of Kcnq3 decreases the M-current. A) The deactivation protocol in voltage clamp starts by holding the cell at −60 mV before a voltage step to −20 mV (300 ms) and followed by 5 mV incremented steps (−25 to −75 mV, 500 ms). The current measured at during the steady state (ss) was subtracted from the instantaneous (Inst) point. B, C) The sag seen following the voltage step was the deactivation of the M-current and was larger in control animals (B) compared to KCNQ3 KD animals (C). D) Graphed as an I-V plot, the M-current was found to be smaller across the range of voltages in KCNQ3 KD mice (two-way ANOVA, main effect of Group, F(1,440)=81.54, p<0.0001) tested with significant *post hoc* differences between −45 and −25 mV (*p<0.05 and **p<0.001, *Bonferroni*). E) XE 991 is a selective blocker of KCNQ channels. The M-current was strongly inhibited with 10 minutes of 40 *μ*M XE bath application. F) The area under the curve for the deactivation protocol was calculated before and after XE application to calculate the percent block of the current. The XE sensitive current is the M-current with the remaining portion due to other K*+* conductances. There was no difference in the effectiveness of XE between control (62.9±5.5%) and KCNQ3-KD (52.8±4.9%) cells (unpaired *t*-test, *t*_(13_) = 1.360 p>0.05). Error bars indicate SEM.

To further test for an increased excitability of the cells the rheobase was measured in current clamp [27]. Each cell was held in a hyperpolarized state at −80 mV while incremental current steps were applied to the cell **(Fig 3G,** bottom). The amount of injected current required to fire an action potential was noted and compared between AgRP^Cre^ mice injected with either the Cas9-Ctrl or KCNQ3-KD virus. We found that the KCNQ3 KD mice required significantly less injected current to fire action potentials (**Fig 3H**), and this effect was significant even after normalizing to the cell capacitance. These results indicate that deletion of *Kcnq3* increases the excitability of NPY/AgRP neurons.

### CRISPR knockdown of Kcnq3 enhances susceptibility of males to overconsumption

Females were employed for PCR and electrophysiological experiments up to this point, however we suspected males might be more sensitive to KCNQ3 knockdown. Unlike in females, the resting membrane potential was significantly depolarized in KCNQ3 KD cells (Ctrl: −52.6±1.3 mV, n=32; KCNQ3-KD: −43.3±1.7 mV, n=39; Unpaired t-test, t_*(69)*_ = 4.641, p<0.0001). In global KCNQ3 KO mice, associated currents are only significantly diminished in cell types known to sparsely express *Kcnq5* [29]. Whereas fasting does not alter expression of *Kcnq5* in NPY/AgRP neurons, OVX females nearly double their relative expression of *Kcnq5* when E2 is exogenously replaced, regardless of their energy state [27]. Therefore, we evaluated behavior in male AgRP^Cre^ mice. Groups were arranged to balance starting weight and fat mass. Weight was then tracked for five weeks, but no difference was seen over this period (**Fig 4 A & B**). This finding would suggest food consumption is the same between groups. In order to test this, mice were individually housed, and their daily food intake measured and, as suspected, no difference was apparent between control (4.5±0.2 g) and KCNQ3-KD (4.2±0.1 g) mice. However, KCNQ subunit expression responds to the energy state of the animal. Following an overnight fast, relative *Kcnq2 and Kcnq3* expression is reduced by almost half with a consequently diminished M-current and increased food intake [27]. If *Kcnq3* expression is already at a nadir, the elevated consumption could be attenuated as both groups displayed a similar increase in post-fast food intake (**Fig 4C**), perhaps due to a ceiling effect.

**Figure 4:**
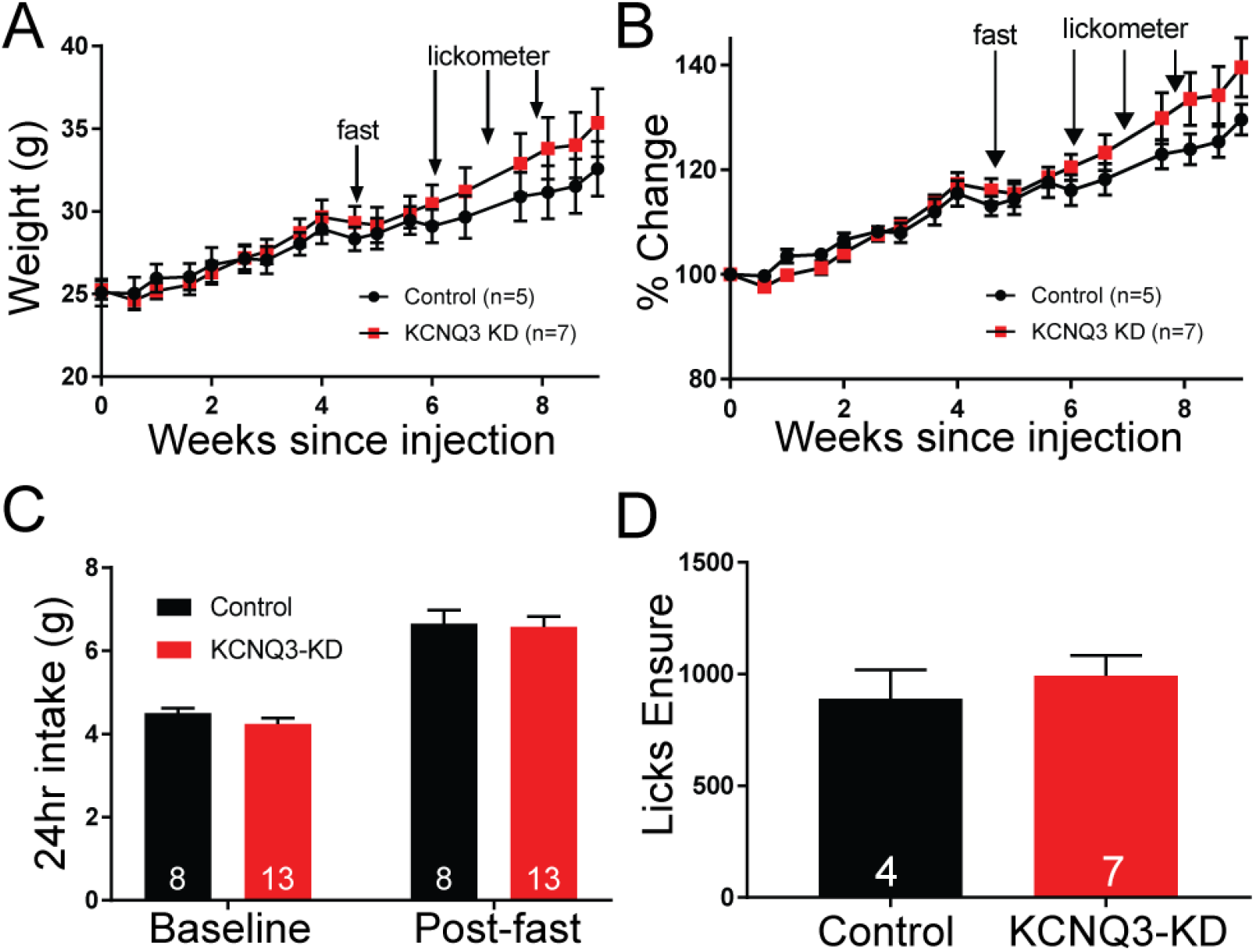
Effects of Kcnq3 knockdown on ingestive behavior. A) AgRP^Cre^ male mice underwent EchoMRI and the measurements were used to balance group assignment. After bilateral ARH injections of either the SaCas9 Ctrl (n=5) or KCNQ3 KD (n=7) virus, mice were regularly weighed. At five weeks post-injection, prior to onset of behavioral testing, the average weight of each group was similar. B) Even after normalization to initial weight, there was no difference in the average percent change between groups. Interestingly, a trend of greater weight gain for KCNQ3 KD mice began to emerge after overnight fast and subsequent exposure to Ensure, a palatable liquid diet (see arrows). C) Mice were individually housed, and their daily intake of standard chow evaluated. Mice were then placed in clean cages with no food for 16 hours. Once food was returned, as expected, mice overconsumed compared to baseline intake. This paradigm is known to decrease expression of both *Kcnq2* and *Kcnq3.* However, there was no difference in baseline or post-fast chow consumption which would suggest a ceiling effect. D) Mice were trained to drink from lickometers containing Ensure, a highly palatable nutritionally complete diet. After four days of training, over a 60 minute access period there were no significant difference between the number of Ensure licks. However, the KCNQ3 KD group did have a higher average than controls (Ctrl: 889±130, n=4; KCNQ3-KD: 993±90, n=7; unpaired *t*-test, *t*93) = 1.198 p>0.05). Error bars indicate SEM.

Palatability is another factor that can drive overconsumption of food [55]. Ensure (Abbott Nutrition) is a nutritionally complete liquid diet often used in feeding and reward studies. Rodents readily consume Ensure with minimal training, even increasing their daily energy intake in a sustained manner [56]. Therefore, ingestive behavior was also measured using a short-term access paradigm. Fed mice during their relative light phase were placed in an operant chamber (Med Associates) equipped with dual lickometers for thirty minutes on four training days. One bottle contained tap water and the other vanilla Ensure (36 g in 100 ml tap water). On the final testing day over sixty minute, while the KCNQ3 KD group did register more licks on average than the control, this difference was not significant (**Fig 4D**).

Finally, NPY/AgRP activity is not just associated with ingestive behavior. Brain infusion of NPY agonists [57, 58] and manipulations that increase NPY/AgRP activity [59, 60] tend to produce anxiolytic effects. The Open Field Test (OFT, **Fig 5A-B**) is a well-established assay for comparing differences in anxiety between groups [61, 62] based on time spent in the center (**Fig 5C**) and frequency of center crossing (**Fig 5D**). Since the surgical control (AAV-mCh) and control vector (SaCas9-Ctrl) injected mice express the same levels of *Kcnq3* mRNA (**Fig 2E**), we used AAV-mCh injected mice for these OFT studies. Interestingly, there was a clear behavioral difference between groups. KCNQ3 KD mice were less anxious, spending more time in the center. Though not as useful as home cage observations, total distance traveled and average velocity (**Fig 5E-F**) provide insight into energy expenditure [63, 64]. Once again, KCNQ3 mice were different than control animals with significantly lower distance and velocity. At the end of the behavioral studies, the mice were euthanized, and the quality of injections confirmed before subjects were included in final analysis. Of the eight KCNQ3-KD animals, four had bilateral injections, three had unilateral injections, and one had no apparent labeling. Those mice with better injections tended to show a more robust behavioral phenotype. For example, decreased distance and velocity were correlated with injection quality (r^2^=0.6, p<0.05, n=7). Together these findings represent subtle, but potentially significant alterations in energy balance. While KCNQ3 animals do not display differences in consumption of standard chow, KCNQ3-KD mice began to accelerate their weight gain following overnight fast and access to palatable food. Coupled with indications of diminished physical activity, animals with KCNQ3 KD specifically in NPY/AgRP neurons may be more susceptible to developing obesity [65, 66].

**Figure 5:**
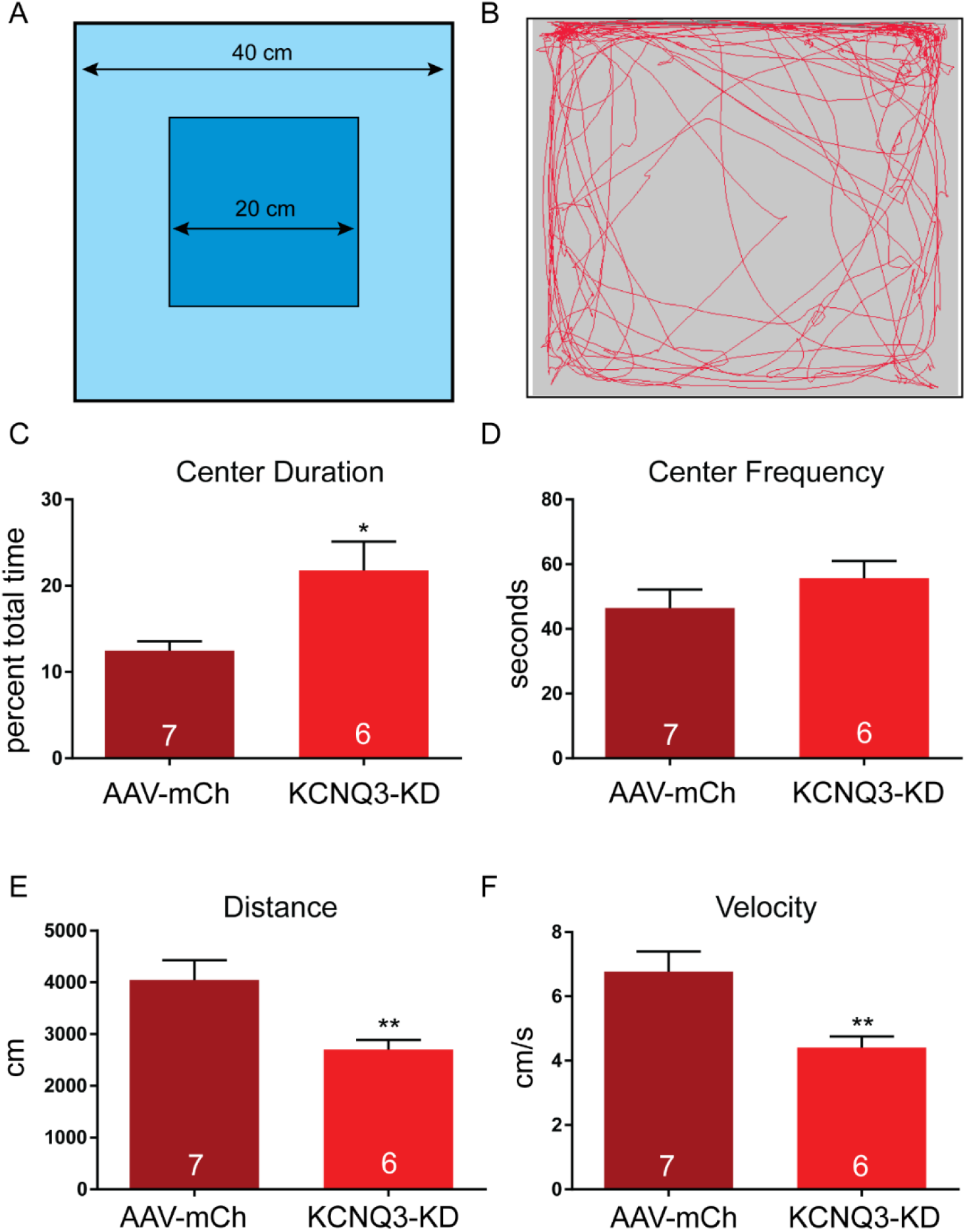
Consequence of Kcnq3 knockdown on open field behavior. A) Open field test (OFT) was done in opaque white boxes 40 cm x 40 cm over 10 min. B) Movement was tracked using video analysis by Ethovision (Noldus). C) KCNQ3 KD increased time spent in the center of during the OFT (Control 12.5±1.1% vs KCNQ3 KD 21.8±3.3%, Unpaired t-test, *t*_(11)_ = 2.851, p<0.01)), indicating an anxiolytic effect. D) The frequency of crossing between the center and periphery was the same between groups (Control 46.4±5.7 vs KCNQ3 KD 55.7±5.3, unpaired *t*-test, *t*(6) = 1.353, p>0.05). E) With the center duration higher, but center frequency the same, it is not surprising that the KCNQ3 animals exhibited lower total distance traveled (Control 4046±384 cm vs KCNQ3 KD 2703±185 cm; unpaired t-test, *t*_(12)_ = 3.147, p<0.01)) and F) lower velocity (Control 6.8±0.6 cm/s vs KCNQ3 KD 4.4±0.8 cm/s; unpaired t-test, *t*_(12_) = 3.147, p<0.01). Statistical analysis was done using t-tests. Error bars indicate SEM.

## Discussion

Using a newly developed CRISPR strategy, we have shown that this single vector approach reduced the expression of *Kcnq3* in NPY/AgRP neurons, resulting in molecular, cellular, and behavioral changes. While this approach was previously validated [41], we have extended this finding using a stringent method of harvesting healthy cells paired with a quantitative measure of relative gene expression [53]. Furthermore, we have confirmed that this change was not due to the surgery, SaCas9-alone, or fluorophore. The single vector SaCas9 system enables a stable (at least 10 weeks) region and cell specific knockdown of a gene in adult animals, avoiding potential developmental compensation or fetal lethality. One limitation of this approach stems from the random nature of the indel. However, Sanger sequencing found only 12% of GFP+ cells retained a wildtype sequence (**Fig 1E**). Most cells exhibited an insertion or deletion that is likely to have formed a frameshifted mRNA [41]. The current construct has the PAM and sgRNA well over 50 nt from an exon junction in the 5’ to 3’ prime direction, which would support rapid non-sense mediated decay of the mutated transcript [67]. Also, the premature stop codon would be generated at the start of the sequence before the first exon junction complex, meaning the decay process is likely efficient [68]. As we selected for fluorescent cells, the remaining transcript detected (**Fig 2B**) is probably edited, non-translatable *Kcnq3* transcripts resistant to nonsense mediated decay pathways [69, 70]. A final innovation is the targeting of ion channel subunits such as *Kcnn3* in studies of Hunker *et al.* [41] and *Kcnq3* in the present study. Unlike the suprathreshold stimulus resulting from optogenetic or chemogenetic stimulation [6, 7], CRISPR deletion of *Kcnq3* increased the overall excitability of NPY/AgRP neurons without directly driving action potentials, but still produced a behavioral phenotype. Therefore, this is a powerful tool for revealing the functional significance of channels in a specific neuronal population in adult animals.

The ability to target subunits is a particularly exciting strategy. Many, if not most, receptors and channels, lack subunit selective agonists and antagonists. The “gold standard” antagonists for studying the M-current, XE991 and linopiridine, are broadly effective. The small molecule UCL2077 [71] does exhibit some subtype selectivity, strongly blocking KCNQ1-2, slightly inhibiting KCNQ3-4, and enhancing KCNQ5 currents [72]. However, heteromeric permutations complicate the pharmacological manipulations. For example, while KCNQ subunits present differing sensitivity to tetraethylammonium, a non-selective K^+^ channel blocker [73], the most common heteromeric complexes are insensitive to the compound. Therefore, other than in heterologous expression systems, pharmacologically untangling the naturally occurring subpopulations of KCNQ channels is a daunting task. Here we have found that deletion of *Kcnq3* attenuates the magnitude of the M-current, but that the subunit is clearly not obligatory. In fact, the relative expression of *Kcnq3* is less than half of *Kcnq2* (**Fig 2D**). This would suggest that most wildtype channels are KCNQ2 homomers or KCNQ2/5 heteromers. Future experiments could target the other subunits. One would expect that in NPY/AgRP neurons deleting *Kcnq2* would produce more robust behavioral alterations and deletion of *Kcnq5* could affect presentation of sex differences. Furthermore, the strategy we have outlined can easily be adapted to study other subunits and cell types.

### The M-current controls resting membrane potential and is suppressed during fasting in NPY neurons

Arcuate NPY/AgRP neurons are involved in the control of food intake and are affected by multiple central and peripheral signals that reflect energy states such as satiety and fasting (reviewed in [11, 74, 75]. In fed animals, NPY/AgRP neurons are less active and exhibit reduced firing activity [13], whereas NPY expression and release is increased during periods of food restriction or deprivation [76–78]. In our previous studies we demonstrated that NPY/AgRP neurons in fed animals were relatively quiescent and more hyperpolarized compared to the fasted state [27]. The depolarized state reported in fasted mice is probably due to a combination of pre-synaptic inputs, changes in peripheral hormonal signals such as leptin [79, 80] and modulation of endogenous potassium conductances like K_ATP_ channels [81, 82] and KCNQ channels [27].

Most arcuate NPY/AgRP neurons express multiple KCNQ subunits and exhibit a robust M-current. One of the primary functions of the M-current is to control action potential firing, spike-frequency adaptation, depolarizing afterpotentials and afterhyperpolarization (AHP) currents [26, 29, 83, 84]. Previously we examined the effects of XE991, a selective blocker of KCNQ channels, on the excitability of NPY neurons from fed males. We found that by blocking the M-current, quiescent NPY neurons became depolarized and started firing continuously with a concomitant reduction in the rheobase current [27]. However, action potential firing did not occur in the presence of glutamatergic and GABAergic blockers although the membrane potential was similarly depolarized. This data suggested that the M-current in hypothalamic NPY/AgRP neurons is important for the control of the resting membrane potential. Although the M-current does have a role in action potential generation in other neuronal cell types (*e.g.* hippocampal CA1 pyramidal neurons) [85, 86], in arcuate NPY/AgRP neurons action potential generation appears to be dependent on the presynaptic inputs (glutamatergic) [87]. Therefore, any alteration in M-current activity could significantly affect the excitability of NPY/AgRP neurons. For example, fasting greatly reduces the M-current while increasing the neuronal excitability of the NPY/AgRP neurons [27]. This effect on M-current activity is partially restored by a 24-hour re-feeding, but not by a shorter 2-hour re-feeding. The suppression of the M-current by fasting may involve peripheral signals such as serum leptin concentrations since leptin does affect Kv2.1 channel activity in NPY/AgRP neurons [82], but it is not known at this time if it affects KCNQ2/3 or the M-current.

### The role of the M-current in the control of food intake by 17β-estradiol

E2 is known to reduce NPY/AgRP expression in OVX females [17, 18, 88] and during proestrus when serum E2 levels are elevated [89]. Short- and long-term E2 replacement reduces NPY mRNA expression and protein immunoreactivity in the arcuate nucleus [17, 18], and long-term (18 days) E2 treatment reduces the *in vitro* release of NPY in the paraventricular nucleus [88]. The suppression of NPY mRNA expression in intact females is associated with a reduction in food intake and body weight, and E2 attenuates food intake during re-feeding (2 hour) after fasting, indicating that the interplay between fasting and E2 is complex [89]. E2 replacement also attenuates the increase in food intake following NPY infusion into the lateral ventricle [90]. These data suggest that the anorectic effects of E2 involve the suppression of NPY expression and neuronal function, which may involve the potentiation of the M-current [27].

NPY/AgRP neurons are significantly hyperpolarized in the E2-versus oil-treated, ovariectomized females, in part, because of the potentiation of the M-current [27]. This potentiation could be due to the KCNQ5 subunit expression. Furthermore, the KCNQ5 subunit contributes to the medium and slow AHP current [31] in hippocampal CA3 neurons, which may also be altered in the E2-treated NPY/AgRP neurons to reduce firing frequency. Ultimately, a reduction in activity would lead to less NPY/AgRP release and a subsequent attenuation in feeding activity. However, fasting abrogates the effects of E2-treatment on M-current activity [27]. This fasting effect is due, in part, to reduction of KCNQ2/3 expression in the neurons resulting in a significant depolarization of the membrane potential, similar to the fasted males. The effects of fasting overwhelmed the effects of E2 (elevated KCNQ5 expression) at the neuronal level via the reduction of the other KCNQ subunits, especially KCNQ3. This is reflected at the behavioral level where estradiol benzoate-treatment does not cause a difference in refeeding compared to oil treatment until three days post-fast [91]. Co-expression of KCNQ3 with KCNQ5 in heterologous cell systems increases current amplitudes 2-3 fold over KCNQ5 homo-multimers and also produces currents with similar amplitudes to the native M-current in hippocampal neurons [31, 36]. Thus, the more robust M-current measured in fed, E2-treated NPY neurons may be due to a greater association of KCNQ3/5 while the effects of fasting reduces this association via a reduction in KCNQ3-containing channels [27].

### Behavioral Consequences of KCNQ3 Knockdown in NPY/AgRP Neurons

NPY/AgRP neurons are critical to the homeostatic regulation of energy balance. In order to fulfill this role, they are able to detect changes in energy reserves [92–95] and drastically alter their connectivity and excitability within hours [79]. The M-current is one mechanism thought to underlie this rapid adaptability [27]. At the start of these studies it was uncertain whether attenuation of the M-current would be sufficient to promote weight gain. Suprathreshold activation, as seen with opto- and chemo-genetic approaches, drives rapid and robust food consumption [6, 7] that would lead to weight gain. Yet, in the present study neither daily consumption of standard chow nor weight gain was different between groups, despite elevated neuronal excitability as evidenced by the change in the rheobase. This led us to hypothesize that challenges might be necessary to draw out an ingestive phenotype. First, we measured intake following overnight fast access and found no difference. Next, mice were given brief access to a vanilla Ensure, since *in vivo* studies have shown that the palatability affects the degree to which NPY/AgRP neurons respond to food presentation [96]. However, neither measure of consumption were significantly different between groups. Interestingly, the group weights and change in weight began to diverge following behavioral experiments. This might suggest that an impaired M-current has a limited effect on food intake; however, dietary challenges (fasting and sugary foods) or age may trigger a KCNQ3 KD phenotype that leads to obesity.

NPY/AgRP neurons make extensive projections throughout the brain to regions not directly involved in food intake [97]. The influence of increased NPY/AgRP activity, therefore, could also manifest as alterations in related, but non-ingestive behaviors [98]. Chronic, chemogenetic activation of NPY/AgRP neurons causes both hyperphagia and reduced physical activity [7]. Other studies have more subtly altered NPY/AgRP neuronal activity without resorting to activation or ablation. Sirt1 is NAD^+^-dependent deacetylase [99] that is expressed in the hypothalamus [100] and is associated with negative energy balance [100–102]. Pharmacological inhibition [103] or knockout of Sirt1 [59] increases NPY/AgRP excitability to reveal increased exploration and greater responses to novelty and cocaine. The open field test represents a simple assay to measure anxiety and, indirectly, energy expenditure. Previously, NPY/AgRP neuronal activation was found to diminish anxiety-related behaviors [60], though this could be due to lowered risk aversion rather than an anxiolytic effect [104]. Lacking available food, the animal may instead display displacement behaviors rather than continue foraging [60]. Therefore, the decreased distance traveled in KCNQ3 KD mice could reflect lower levels of physical activity (distance traveled), change in anxiety, or displacement behaviors in the absence of food. It should be noted that we noted this decrease during the relative light phase when the animal would normally be at rest. Future experiments could examine if KCNQ3 KD affects energy expenditure or circadian rhythms. In summary, our data demonstrates that the M-current is an important cellular mechanism by which NPY/AgRP activity is regulated in response to the steroid and energy state of the animal. Significantly, loss of this channel subunit can have clear effects on specific behaviors that could predispose mammals to developing obesity.

## Acknowledgements

This research was supported by United States Public Health Grants DK 68098 (MJK and OKR), NS 43330 (OKR), NS 38809 (MJK) and MH 104450 (LZ).

